# Rhizobacteria inoculation of plants for reducing insect herbivores: A meta-analysis on insect behaviour and fitness

**DOI:** 10.1101/2024.04.18.590063

**Authors:** Sharon E Zytynska, Megan Parker, Oriana Sanchez-Mahecha

## Abstract

Microbial communities in the plant rhizosphere—the soil region closely associated with plant roots— play critical roles in shaping plant growth, defence, fitness, and ecosystem processes. Inoculation of plants with specific rhizobacteria strains has shown promising potential for increasing crop yields. Rhizobacteria can also induce plant defences, resulting in reduced insect growth and reproduction, and can manipulate plant biochemistry to alter insect host-choice and recruit natural enemies of the insects. We present a meta-analysis examining the impact of rhizobacteria inoculation of plants on insect fitness and behaviour. Our findings indicate that rhizobacteria inoculation generally reduces herbivore fitness and host choice behaviours. However, effect sizes varied significantly depending on type of herbivore (chewing versus sucking), plant host, and rhizobacterial species. *Bacillus* spp. showed stronger effects than the commonly studied *Pseudomonas* spp. Rhizobacteria notably reduced traits such as host choice, leaf consumption, survival, and reproduction of chewing herbivores, while primarily impacting sucking herbivores by reducing reproduction. Single-strain inoculants tended to perform better, especially for sucking herbivores, suggesting potential strain incompatibility issues with multi-strain inoculants. Furthermore, field trials showed less impact on insect fitness reduction compared to experiments under controlled conditions, possibly due to soil diversity and environmental factors affecting inoculant persistence. Despite, very limited experimental data, studies observed that rhizobacteria inoculation of plants can attract parasitoid wasps and predators to the plants. These results underscore the need for considering broader environmental interactions when developing effective rhizobacteria-based pest management strategies. Understanding specific and generalist rhizosphere interactions can aid in developing synthetic microbial communities with broad protective functions across various plants and environments.

## Introduction

The rhizosphere is the narrow region of soil directly influenced by plant roots, hosting a dynamic and diverse microbial community with up to 10^11^ microbial cells per gram root and more than 30,000 prokaryotic species (Berendsen, Pieterse & Bakker 2012). This zone plays a vital role in sustaining plant health by fostering interactions between roots and beneficial microorganisms. Plants influence the rhizosphere microbiome by releasing root exudates and altering soil conditions, selectively recruiting distinct microbial communities tailored to a plant’s unique physiological and biochemical traits (Berendsen, Pieterse & Bakker 2012; Trivedi *et al*. 2020). Recruitment of rhizosphere bacteria can occur over short-time frames, enabling a plant to efficiently respond to changes in the environment, e.g. from drought or nutrient stress (Bakker *et al*. 2013). Recent syntheses examining the effect of rhizobacterial inoculation on plant responses to drought (Rubin, van Groenigen & Hungate 2017; Zhao *et al*. 2022), temperature (Zhang *et al*. 2023), salt (Pan *et al*. 2019), phosphate (De Zutter *et al*. 2022) and across different experimental factors (Zeffa *et al*. 2020; Li *et al*. 2022; da Silva *et al*. 2024) have demonstrated the strong and broad impact that different rhizobacteria can have on plant fitness and yield.

Plants can also respond to herbivory through recruitment of different rhizobacteria communities, that enhance plant defences responses, thereby decreasing insect population growth rates (Pineda *et al*. 2010). Even before herbivory, plants can be primed to respond more quickly upon insect feeding through application of various plant hormones but also inoculation with defence-inducing rhizobacteria (Pieterse *et al*. 2013; Pieterse *et al*. 2014; Selosse, Bessis & Pozo 2014). Priming of a plant involves structural changes in the histones that fold DNA, leading to a quicker initiation of RNA transcription and faster synthesis of defence-related compounds. The mechanisms underlying this vary across plant and herbivore species, with variable induction of defences by different rhizobacteria.

Plant responses to chewing insects, e.g. caterpillars, differ from responses to sucking insects, e.g. aphids or whitefly (Wu & Baldwin 2010; Schuman & Baldwin 2016). The complex interplay between plant responses and the type of herbivory suggests that the effectiveness of rhizobacteria is context-dependent (Blubaugh *et al*. 2018; Friman *et al*. 2021). Chewing insects often trigger robust wound-response pathways, activating jasmonic acid signalling, whereas sucking insects primarily induce salicylic acid pathways, leading to different plant defence mechanisms (Walling 2000). This divergence in defence strategies implies that rhizobacteria-mediated protection may be specific to the type of herbivore involved, highlighting the importance of understanding these relationships to optimize biological control strategies.

Here we present a meta-analysis to synthesise published data measuring effects of experimental bacterial inoculation of plants on insect behaviour and fitness, comparing chewing and sucking herbivorous insects. In addition to defence induction, microbial inoculation and herbivory can also trigger changes in plant volatiles that can recruit natural enemies of the herbivores. While Herbivore-Induced Plant Volatiles (HIPVs) in plants are well-known to attract natural enemies (Turlings & Erb 2018), the consequences of microbial-induced plant volatiles on herbivore or natural enemy behaviour is relatively unknown. We also examined parasitoid wasp and predator behaviour in response to bacterial-inoculation of plants. This meta-analysis aims to describe general effects of experimental microbial inoculation of plants on insect behaviour and fitness, with a focus on rhizosphere bacteria effects on chewing and sucking insects.

### Our main hypotheses are

1. The effect of microbial inoculation on insect responses varies across insect feeding groups, insect family and host plant family
2. The type of inoculation (single or community) as well as the main bacterial genus in the inoculant will impact insect responses
3. The experimental environment (controlled pot experiments vs field trials) will alter the strength of response of the insect

## Methods

We used two broad and inclusive search approaches to maximise identification of papers for inclusion. The first search was compiled in ISI Web of Science up to September 2023 using the following keywords combinations: (Rhizo* AND aphid*), (Rhizo* AND Whitefly*), (Rhizo* AND cater*), (Rhizo* AND lepido*), (Rhizo* AND herbivor*), resulting in 428 papers. We used separate terms for aphid, whitefly, caterpillars, and lepidoptera as these are well-known species studied in this area, and the term herbivore was used to encompass a wider range. The second search used three sub-searches using terms 1. (“Insect*” AND (“volatile*” OR “terpen*” OR “sesquiterp*” OR “floral volatile*” OR “leaf volatile*”) AND (“microb*” OR “bacteri*” OR “rhizo*” OR “fungi*”) AND (“behav*” OR “attract*” OR “repel*” OR “pref*” OR “oviposit*” OR “feed*”), 2. (“Insect*” OR “herbiv*” OR “behav*” OR “parsit*”) AND (“volatile*” OR “terpen*” OR “sesquiterp*” OR “floral volatile*” OR “leaf volatile*”) AND (“microb*” OR “bacteri*” OR “fungi*”) AND (“plant*”) AND (“behav*” OR “attract*” OR “repel*” OR “pref*”), 3. (“Insect*” OR “herbiv*” OR “behave*” OR “parsit*”) AND (“volatile*” OR “terpen*” OR “sesquiterp*” OR “floral volatile*” OR “leaf volatile*” OR “secondary metabol*”) AND (“microb*” OR “bacteri*” OR “fungi*”) AND (“behav*” OR “attract*” OR “repel*” OR “pref*”) resulting in 1615 papers. Following assessment of title and abstract, we identified 180 potential papers of interest, from which 70 provided suitable data for inclusion in the meta-analysis (Fig. 1).

**Fig. 1.**
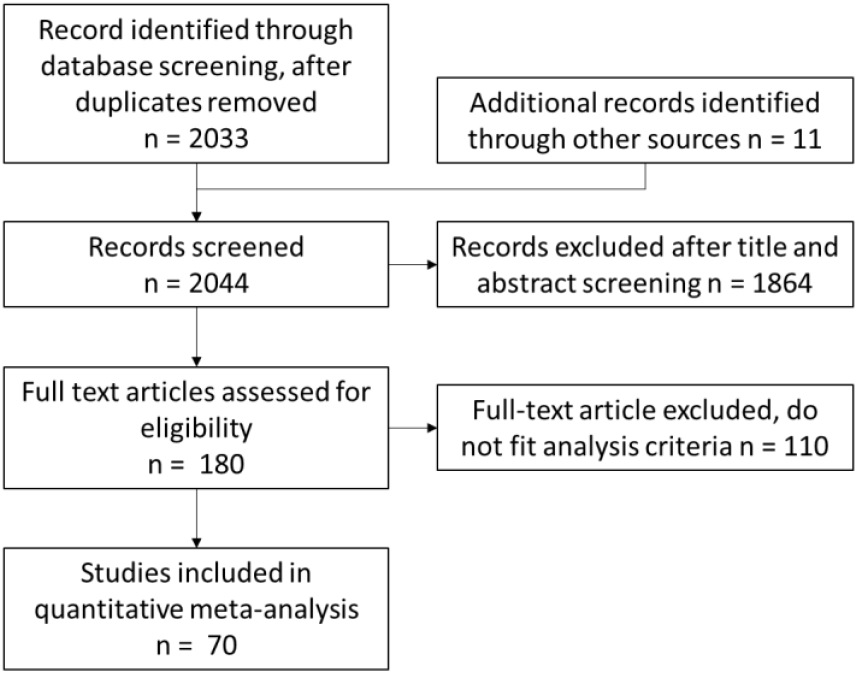
Summary of selection process for final papers for inclusion in the meta-analysis, as based on PRISMA guidelines

We screened the studies using the following inclusion criteria: (1) data must come from experimental or field studies that tested the bacteria effect on insects comparing a bacteria-treated with control untreated plants; (2) data are presented to enable extraction of mean values, a measure of variance, and sample size; (3) measured variables are related to insect behaviour (host choice or consumption rate) or fitness (insect survival, development time, body size, fecundity, or lifespan). We extracted the mean, variance, and sample size across the treated and untreated groups. We also extracted experimental information on bacteria inoculum (single strain or multi-strain community), microbial genus/species (multi-genus noted for mixed-genera community inoculations), insect feeding type (chewing herbivore, sucking herbivore, parasitoid wasps or predator), insect family/species, plant family/species, inoculation method (soil, seed, root, leaf), and study environment (field, greenhouse, or growth chamber). Following in-depth screening of 180 papers, 44 papers were removed due to not containing primary data, 66 removed due to lack of informative data across treatments and/or controls, resulting in a final data set of 344 data points extracted from 70 papers published between 1997 and 2023 fulfilling all the inclusion criteria.

In studies with multiple genotypes, strains, or varieties, data were pooled within species due to lack of replication to examine specific effects. We selected only data from the “wildtype” microbial strain where data were presented also on lab-generated mutants. For controlled environment time-series experiments we chose the time that related best to the other studies in the dataset; for example, the number of offspring produced by aphids is commonly measured at day 10 or 14. For field data, we averaged across multiple sampling time points.

### Data analysis

The meta-analysis was performed in RStudio version 2024.04.2 (RStudio Team 2024) using R version 4.3.0 (R Core Team 2024) using the package metaphor (Viechtbauer 2010). Effect sizes were calculated using the standardized mean difference with an unbiased estimate of the sampling variances (SMDH, giving Hedges’ g). This measurement compares the treated plants (inoculated with the bacteria and exposed to the herbivore insect) to the control plants (that were just exposed to the herbivore insect). A priori power analyses (medium effect size d=0.5) were calculated in R following Quintana and Tiebel (2019) for the main insect variables within the analysed data subset. For the power analysis, we used the number of effect sizes from the extracted data (data points), the average number of replicates used to calculate each effect size (control vs treated plant), against a predicted effect size of d = 0.5 (medium) and low heterogeneity (h = 0.33).

The effect of the inoculated bacteria on the insect traits was analysed with a meta-analytic linear mixed effect model (rma.mv). Study was included as a random effect to account for multiple data points across insect, plant, and bacteria species within each study. In each insect subset (sucking and chewing herbivores), insect response was treated as a fixed effect moderator (levels: host choice, feeding, survival, development, body size, fecundity, lifespan) also separated by main rhizobacteria genera (levels: *Bacillus, Pseudomonas*, Other) using a nested analysis approach. Additional models were used to determine the effect across different plant and insect families, as well as across experimental manipulations (controlled pot experiments vs field trials). Importantly, only results are presented when there were at least three data points from three independent studies for the comparisons. We evaluated publication bias in the dataset by assessing the funnel-plot asymmetry and Eggers test. For the data visualisation, figures show both the mean effect size and 95% confidence intervals obtained from the linear mixed effect models output, as well as the individual-level data as a bubble plot with sizes relative to the precision of the estimate (1/SE), where larger bubbles show a higher certainty of the data. *k* corresponds to the number of effect sizes, from *n* number of studies.

## Results

### Effects of rhizobacteria inoculation of plants on associated insects

The herbivore final dataset contained 344 data points from 66 papers, representing 215 data points (38 papers) from chewing herbivores and 129 data points (31 papers) from sucking herbivores, with data on all responses identified (Table 1). Across all measured response traits in all herbivore species, rhizobacteria inoculation of plants had a negative effect on insect herbivores (hedges g = - 0.417, P<0.001, k=344). Yet, the strength and significance of these effects varied between chewing (hedges g = -0.563, P<0.001, k=215) and sucking herbivores (hedges g = -0.242, P=0.209, k=129), as well as across the different response traits measured (test of moderators QM=59.3, df=14, P<0.001; Fig. 2a,b). A large residual heterogeneity in this model (QM=1416.3, df=330, P<0.001) indicated the need to include additional factors in the model (rhizobacteria/plant/insect species or system information) and therefore in following sections we examined chewing and sucking herbivore datasets separately.

**Table 1.** Summary of data points included in meta-analyses, separated by seven behavioural and fitness traits commonly measured, exploring the distribution of data points among experimental factors.

|  | Number of data points (number of articles) |  |  |  |  |  |  |  |
| --- | --- | --- | --- | --- | --- | --- | --- | --- |
|  | host choice | feeding | survival | development | body size | fecundity | lifespan | k |
| Chewing herbivores | 25 (8) | 26 (8) | 24 (6) | 39 (8) | 72 (24) | 19 (7) | 10 (3) | 215 |
| controlled pot | 10 (3) | 25 (7) | 24 (6) | 39 (8) | 72 (24) | 18 (6) | 10 (3) | 198 |
| field | 15 (5) | 1 (1) |  |  |  | 1 (1) |  | 17 |
| single strain | 15 (6) | 21 (7) | 22 (6) | 37 (8) | 61 (23) | 7 (5) | 10 (3) | 173 |
| community | 10 (5) | 5 (2) | 2 (1) | 2 (1) | 11 (3) | 12 (4) |  | 42 |
| Sucking herbivores | 2 (1) | 18 (2) | 3 (1) | 19 (8) | 4 (3) | 73 (26) | 10 (5) | 129 |
| controlled pot | 2 (1) | 18 (2) | 3 (1) | 19 (8) | 4 (3) | 44 (20) | 10 (5) | 100 |
| field |  |  |  |  |  | 29 (7) |  | 29 |
| single strain |  | 18 (2) | 3 (3) | 16 (6) | 4 (3) | 63 (26) | 9 (4) | 113 |
| community | 2 (1) |  |  | 3 (3) |  | 10 (6) | 1 (1) | 16 |
| Natural enemies | 19 (7) | 6 (1) | 1 (1) | 1 (1) |  | 4 (1) |  | 31 |
| controlled pot | 10 (5) | 6 (1) | 1 (1) | 1 (1) |  |  |  | 18 |
| field | 9 (2) |  |  |  |  | 4 (1) |  | 13 |
| single strain | 15 (5) | 6 (1) | 1 (1) | 1 (1) |  | 3 (1) |  | 26 |
| community | 4 (3) |  |  |  |  | 1 (1) |  | 5 |

*Notes: Grouping factors are controlled pot vs field inoculation, and single strain vs community (multi-strain) inoculation. Natural enemies includes* *parasitoid wasps and predators combined. k corresponds to the number of effect sizes; the number of studies is shown in brackets.*

**Fig. 2.**
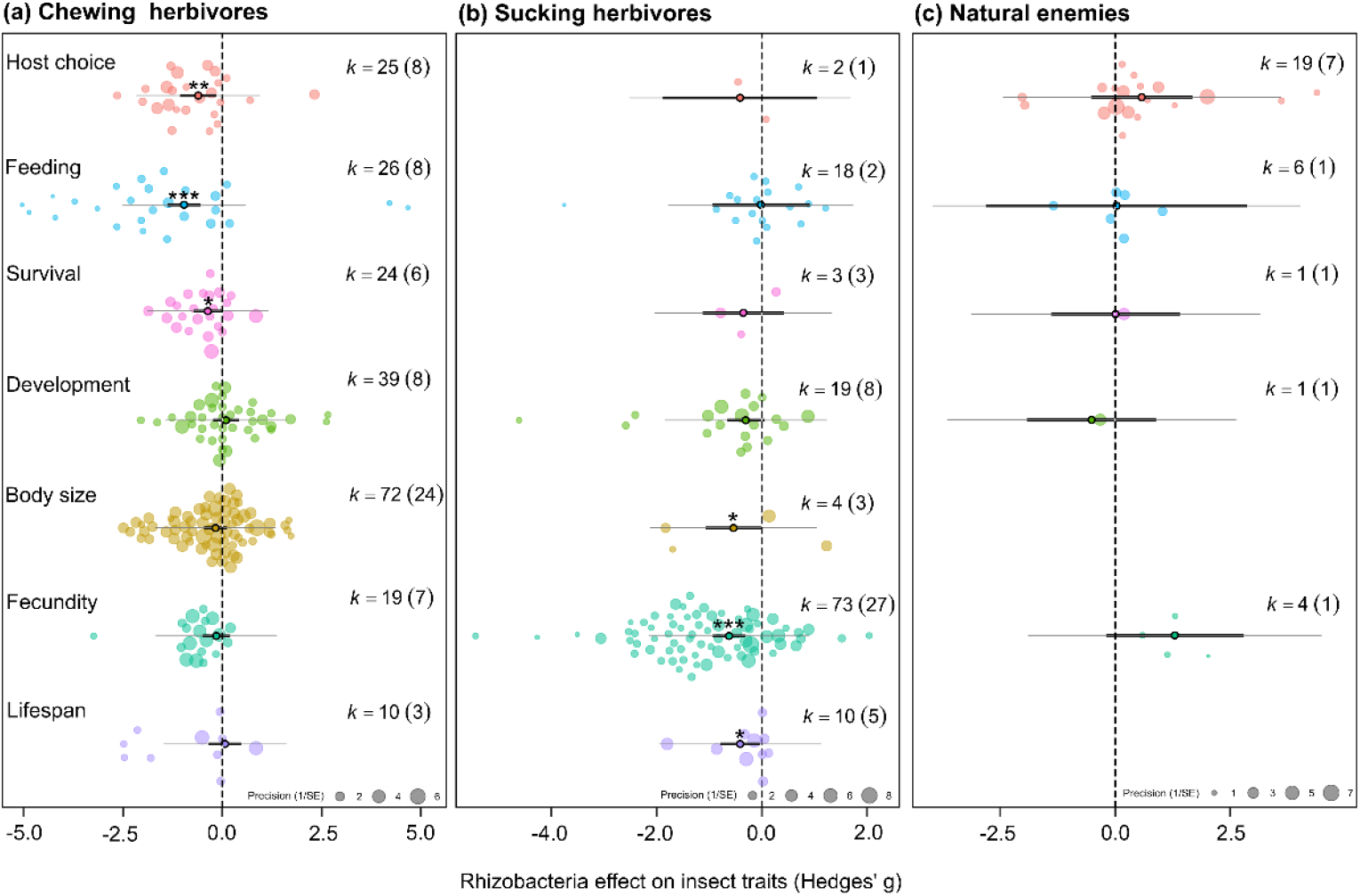
Effect of rhizobacterial inoculation across response traits measured for (a) chewing herbivores, (b) sucking herbivores and (c) natural enemies (parasitoid wasps and predators). Orchard plots showing effect size mean, thick bars—95% confidence intervals, thin bars—95% prediction intervals, and individual effect sizes scaled by their precision (1/SE). *k* corresponds to the number of effect sizes; the number of studies is shown in brackets. Effects are considered significant if the 95% CI does not overlap with zero.

We also extracted 31 data points from eight papers with information on rhizobacterial effects on insect natural enemies (predators and parasitoid, examining variables including host choice, feeding behaviour, and fecundity; Table 1, Fig. 2c). While there were no general significant effects due to a lack of data (hedges g = 0.416, P=0.385, k=31), there is a potential pattern of increased host choice to plant inoculated with non-*Bacillus* or non-*Pseudomonas* rhizobacteria (‘Other’ rhizobacteria: hedges g = 1.61, P=0.050, k=4; *Bacillus*: hedges g = 0.232, P=0.810, k=11; *Pseudomonas*: hedges g = -0.649, P=0.513, k=4). The other rhizobacteria grouping included plants inoculated with *Bradyrhizobium japonicum* and *Delftia acidovorans*.

### Effect of rhizobacteria inoculation on chewing insects

The chewing insects belonged to six different families (Lepidoptera k=178, Chrysomelidae n=19, Anthomyiidae n=15, Cecidomyiidae n=1, Gryllotalpidae n=1, Sciaridae n=1) and the Lepidoptera represented nine genera, including *Spodoptera* sp. (n=48 data points, from 13 papers) and *Plutella xylostella* (n=67 data points, from 4 papers); with a tendency for a small number of papers to examine a broad range of rhizobacteria leading to a loss of independence in these effect estimates.

Effects on chewing insect behaviour were observed across the range of bacterial inoculants leading to overall significant reductions in the number of insects landing on an inoculated host plant (host choice: hedges g = -0.637, P=0.006, n=25, studies=8) with subsequent reduction in leaf consumption (feeding: hedges g = -0.965, P<0.001, n=26, studies=8) (Fig 2a, 3a). There was also a significant negative effect on chewing herbivore survival (hedges g = -0.367, P=0.041, k=24, n=6; Fig. 2a), but this was driven by two studies using *Bacillus* inoculants (hedges g = -0.602, P=0.028, k=11, n=2; Fig. 3a). No significant effect was observed for studies using *Pseudomonas* inoculants (hedges g = -0.131, P=0.656, k=8, n=3; Fig. 3a) or ‘Other’ inoculants (hedges g = -0.150, P=0.586, k=5, n=2; Fig. 3a). Despite body size being the most measured variable, it showed no strong significant reduction when averaged across all rhizobacteria inoculants (hedges g = -0.170, P=0.248, k=72, n=24; Fig. 2a). However, when grouped by rhizobacteria genus, *Bacillus* inoculations of host plants significantly reduced chewing herbivore body size (hedges g = -0.487, P=0.032, k=24, n=6), while *Pseudomonas* and ‘Other’ inoculations (representing nine genera) did not (*Pseudomonas*: hedges g = -0.051, P=0.820, k=39, n=13; Other: hedges g = -0.062, P=0.804, k=9, n=6; Fig. 3a). Insect development time was positively affected by *Bacillus* inoculation of the host plant, representing shorter development time, but only from two studies (hedges g = 0.561, P=0.026, k=19, n=2).

**Fig. 3.**
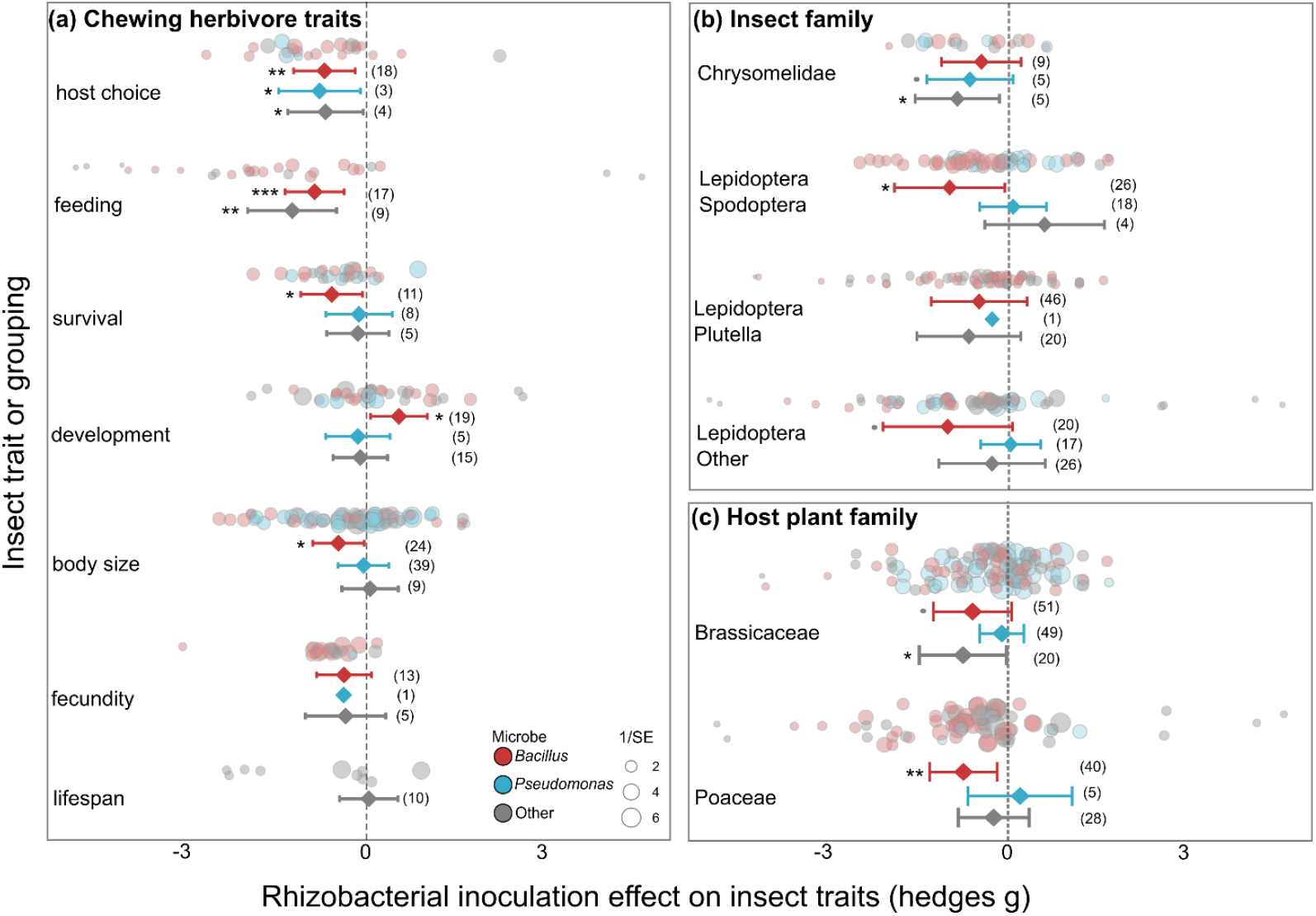
Effect of rhizobacterial inoculation on chewing herbivores, separated by *Bacillus* spp inoculants (red), *Pseudomonas* spp. inoculants (blue), and other rhizobacteria (grey) across (a) response traits (b) main insect families, and within Lepidoptera and (c) host plant family. Effect size mean and 95% confidence intervals from the meta-analysis linear mixed effect model outputs, individual data shown as bubble plots with sizes relative to the precision of the estimate (1/SE), where larger bubbles show a higher certainty of the data. Number of data points (k) shown in brackets.

Of the six chewing herbivore families studied, rhizobacteria inoculation significantly reduced chewing herbivore fitness within Chrysomelidae (hedges g = -0.584, P=0.030, k=19, n =8) and Lepidoptera (hedges g = -0.382, P=0.008, k=178, n =26) across a diverse set of microbial inoculants. Chrysomelidae leaf beetles were negatively affected by non-*Bacillus* bacterial inoculants of their host plants (Other: hedges g = -0.844, P=0.023, k=5, n=4; *Pseudomonas*: hedges g = -0.628, P=0.098, k=5, n=2; *Bacillus:* hedges g = -0.433, P=0.218, k=9, n=4; Fig. 3b). Whereas, within Lepidoptera the strongest effects were from *Bacillus* inoculants (Spodoptera: hedges g = -0.933, P=0.021, k=26, n=3; Other Lepidoptera: hedges g = -1.00, P=0.089, k=20, n=3; Fig. 3b). There were no significant effects on *Plutella* Lepidopterans (Fig. 3b).

Chewing insects were most often studied on Brassicaceae (120/215) or Poaceae (73/215) plants with both Chrysomelidae and Lepidoptera studied within both plant families. *Bacillus* inoculants within both plant families had negative effects on insects (Poaceae: hedges g = -0.818, P=0.006, k=40, n=5; Brassicaceae: hedges g = -0.663, P=0.051, k=51, n=4; Fig. 3c). Other microbial inoculants (including Alcaligenes, Enterobacter, and Kluyvera) also showed significant effects on chewing insects but data were collected from a single paper and thus robustness is low (hedges g = 0.826, P=0.030, k=20, n=1; Fig. 3c). *Pseudomonas* inoculants had no overall effect within Brassicaceae (hedges g = -0.161, P=0.405, k=49, n=12; Fig. 2c) or Poaceae (hedges g = 0.152, P=0.738, k=5, n=3; Fig. 3c).

### Effect of rhizobacteria inoculation on sucking herbivores

The data were dominated by studies on Aphididae (86/129 data points from 8 species of aphid); these included *Brevicoryne brassicae* (n=23), *Acyrthosiphon pisum* (n=18), *Myzus persicae* (n=14), *Rhopalosiphum padi* (n=13), *Lipaphis erysimi* (n=18), *Aphis glycines* (n=5), undescribed aphids (n=4), *Aphis gossypii* (n=3), and *Sitobion avenae* (n=2). The remaining data were from Tetranychidae spider-mites (n=28 on *Tetranychus urticae*), Aleyrodidae whitefly (n=7 from Bemisia tabaci and n=4 from *Aleyrodes proletella)*, Triozidae psyllids (n=3, from *Bactericera cockerelli*), and n=1 from the Cicadellidae leafhopper *Amrasca biguttula*.

Most studies measured sucking herbivore fecundity (number of offspring produced) (73/129) which was significantly reduced by rhizobacteria inoculation of the host plants (hedges g = -0.624, P<0.001, k=73, n=26; Fig 2b). Averaged across all inoculants, we also found significant effects on sucking herbivore lifespan (hedges g = -0.413, P=0.030, k=10, n=5), body size (hedges g = -0.539, P=0.046, k=4, n=3). Similar to the chewing herbivores, the majority of experiments on sucking herbivores used either *Bacillus* (n=71/129) or *Pseudomonas* strains (n=33/129). We found substantial variation in the number of studies and strength of effects across these different inoculants (Fig. 4a). Insect fecundity was significantly reduced by *Bacillus* (hedges g = -0.565, P=0.036, k=42, n=12), *Pseudomonas* spp. (hedges g = -0.559, P=0.033, k=17, n=9), and other inoculant species (hedges g = -0.922, P<0.001, k=14, n=10). A potential negative impact on development time to adult (hedges g = -0.308, P=0.090, k=19, n=8) was driven by a significant negative effect on *Bacillus* inoculated plants (hedges g = - 0.726, P=0.011, k=7, n=3) weakened by a potential positive effects on *Pseudomonas* inoculated plants (hedges g = -0.544, P=0.065, k=7, n=4).

**Fig. 4.**
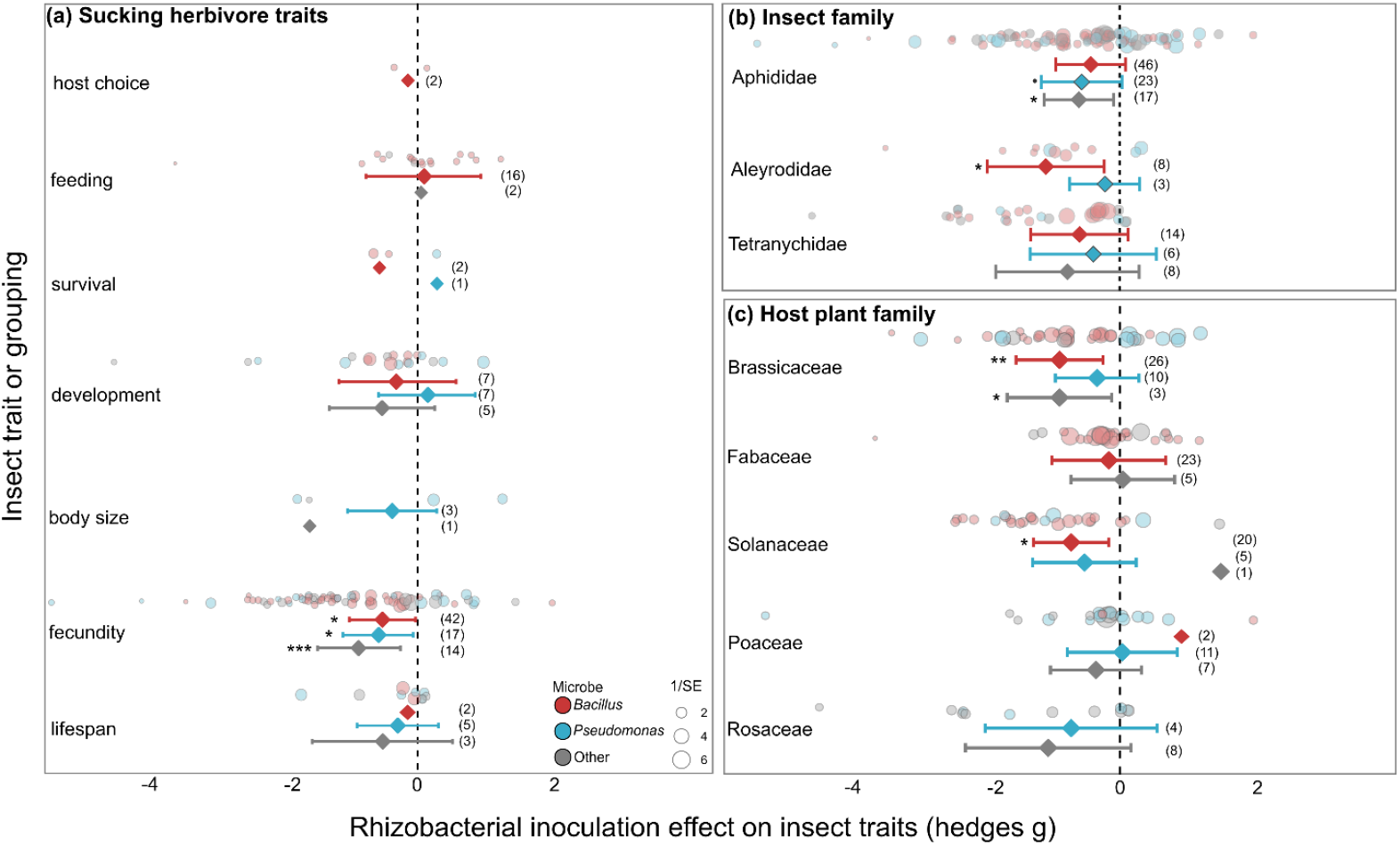
Effect of rhizobacterial inoculation on sucking herbivores, separated by *Bacillus* spp inoculants (red), *Pseudomonas* spp. inoculants (blue), and other rhizobacteria (grey) across (a) response traits (b) main insect families, and (c) host plant family. Effect size mean and 95% confidence intervals from the meta-analysis linear mixed effect model outputs, individual data shown as bubble plots with sizes relative to the precision of the estimate (1/SE), where larger bubbles show a higher certainty of the data. Number of data points (k) shown in brackets.

The three insect families, with sufficient data for analysis, were all negatively impacted by rhizobacterial inoculation of the host plants (Aphididae: hedges g = -0.469, P=0.005, k=86, n=24; Aleyrodidae: hedges g = -0.767, P=0.027, k=11, n=5; Tetranychidae: hedges g = -0.607, P=0.059, k=28, n=3). In particular, Aleyrodidae insects (whitefly) were significantly impacted by *Bacillus* inoculants (hedges g = -1.086, P=0.016, k=8, n=3; Fig 4b), while Aphididae (aphids) were most negatively affected by non-*Bacillus* inoculants (‘Other’: hedges g = -0.589, P=0.027, k=17, n=10; *Pseudomonas*: hedges g = -0.514, P=0.097, k=23, n=8; Fig 4b). For aphid studies, the other inoculants included *Rhizobium* sp, *Bradyrhizobium* sp, *Sinorhizobium* sp, and an *Acidovorax* sp.

Sucking insects were studied across five different plant families (Fig 4c). We found significant negative effects of rhizobacterial inoculation on sucking insects feeding on plants within Brassicaceae (hedges g = -0.755, P<0.006, k=39, n=10), driven by effects of *Bacillus* spp (hedges g = -0.931, P=0.006, k=26, n=4) and other inoculants (hedges g = -0.933, P=0.022, k=3, n=2), but not *Pseudomonas* (hedges g = -0.355, P=0.277, k=10, n=4) (Fig 4c). *Bacillus* spp inoculants also significantly impacted sucking insects on Solanaceae plants (hedges g = -0.753, P=0.011, k=20, n=5; Fig. 4c).

### Effect of rhizobacteria community inoculations (multi-*Bacillus* and multi-genus inoculants)

Chewing herbivores were slightly more negatively affected when feeding on plants inoculated with multiple rhizobacteria (hedges g = -0.383, P=0.043, k=42, n=9) than single inoculants (hedges g = - 0.273, P=0.051, k=173, n=34) (community vs single: QM= 5.04, df=2, P=0.081). This difference was driven by the majority of community inoculations using multiple *Bacillus* strains (hedges g = -0.653, P=0.002, k=39, n=7) rather than multi-genus communities, for which there was insufficient information for a robust analysis (hedges g = -0.511, P=0.330, k=3, n=2). The effect size of multi-strain *Bacillus* inoculants on chewing herbivores was comparable to single-strain *Bacillus* inoculations (hedges g = -0.598, P=0.002, k=63, n=10; Fig 5a). Single-strain inoculants from *Pseudomonas* had only minimal overall effect on chewing herbivores (hedges g = -0.296, P=0.064, k=56, n=16; Fig 5a), an no multi-strain *Pseudomonas* communities were tested.

**Fig. 5.**
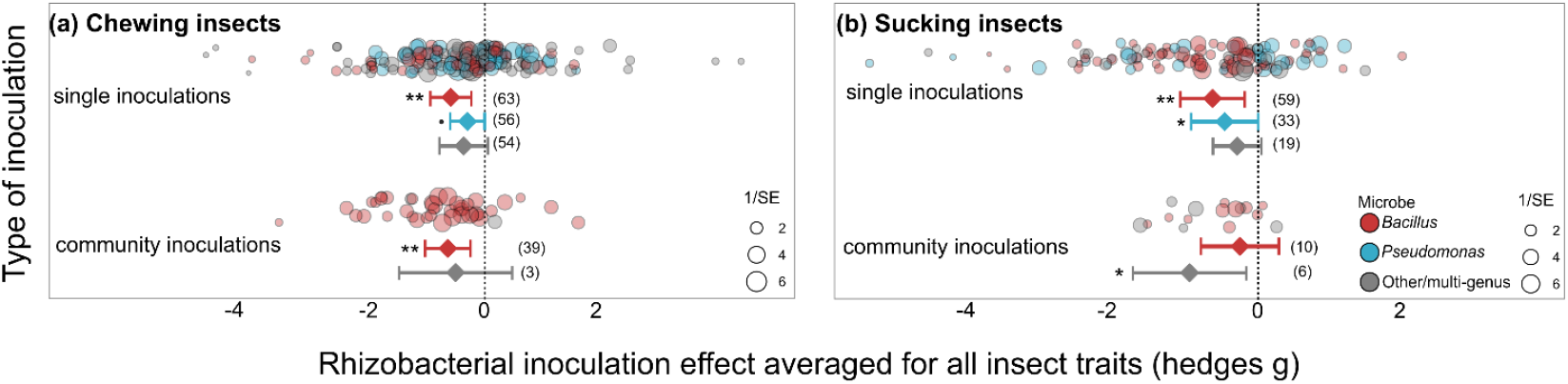
Effect of rhizobacterial inoculation using single and community inoculations for (a) chewing or (b) sucking herbivores. Effect size mean and 95% confidence intervals from the meta-analysis linear mixed effect model outputs, individual data shown as bubble plots with sizes relative to the precision of the estimate (1/SE), where larger bubbles show a higher certainty of the data. Number of data points (k) shown in brackets.

**Fig. 6.**
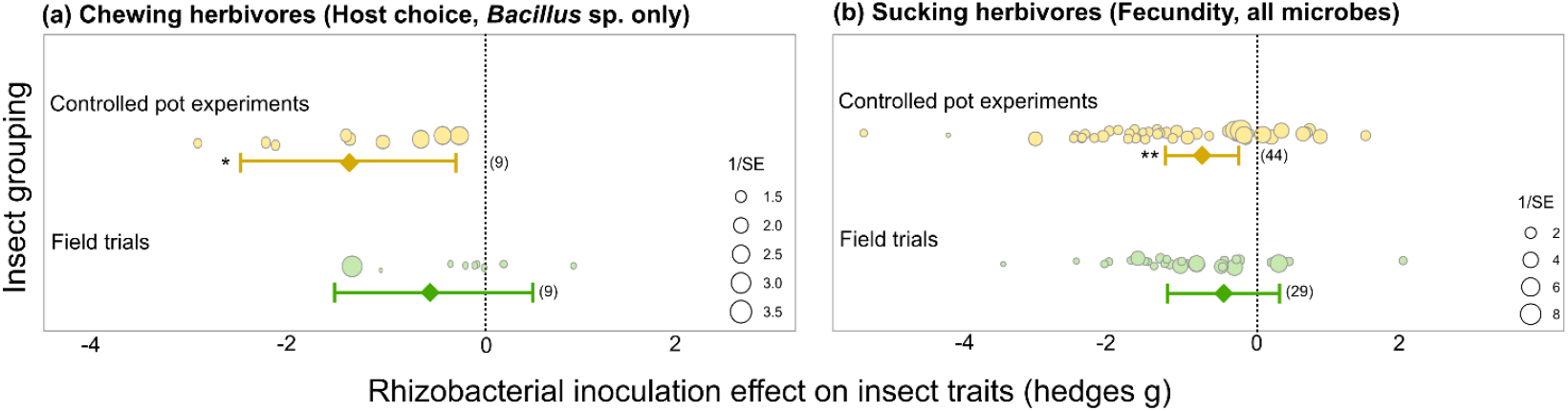
Effect of rhizobacterial inoculation comparing controlled pot and field trials for (a) chewing herbivore using host-choice and *Bacillus* sp only due to lack of data for others, and (a) sucking herbivore fecundity combining all microbes studied. Effect size mean and 95% confidence intervals from the meta-analysis linear mixed effect model outputs, individual data shown as bubble plots with sizes relative to the precision of the estimate (1/SE), where larger bubbles show a higher certainty of the data. Number of data points (k) shown in brackets.

In contrast, sucking insects were more negatively affected by single (hedges g = -0.552, P=0.003, k=113, n=28) than less-studied community inoculations (hedges g = -0.348, P=0.094, k=16, n=8). Community inoculations were dominated by those using multi-strain *Bacillus* inoculants that, surprisingly, had no significant effect on insect fitness (hedges g = -0.236, P=0.393, k=10, n=5) despite single-strain *Bacillus* inoculants having a strong effect (hedges g = -0.624, P=0.007, k=59, n=10). Of particular note, multi-genus inoculants were able to reduce sucking insect fitness yet examined across very few studies (hedges g = -1.016, P=0.010, k=6, n=3). Multi-genus inoculants contained various combinations of *Bradyrhizobium* spp, *Azospirillium* spp, *Delftia* spp, *Azotobacter* spp, as well as *Bacillus* spp and *Pseudomonas* spp.

### Effect of controlled pot experiments compared to field trials

For chewing herbivores, only the host-choice of insects to *Bacillus*-inoculated plants could be analysed to compare field choice and controlled pot experiments since other variables were only measured in controlled pot experiments. Similarly, for sucking insects only insect population sizes were measured (fecundity) across both environments, with multiple bacterial inoculants (minimum of three data points per microbe grouping per environment). We found that for these variables that the significant effect of rhizobacteria in controlled pot experiments (chewing: hedges g = -1.244, P=0.023, k=9, n=3; sucking: hedges g = -0.748, P=0.004, k=44, n=20) was lost in field trials (chewing: hedges g = -0.646, P=0.235, k=9, n=5; sucking: hedges g = -0.449, P=0.256, k=29, n=8).

## Discussion

Overall, inoculation of host plants with rhizobacteria reduces insect fitness and alters behaviour to further reduce colonisation (host choice) and leaf consumption (feeding). We also observed potential for rhizobacteria inoculation to increase host choice of natural enemies of herbivore insects, increasing top-down predator control. This supports continued efforts to understand how rhizobacteria boost plant resistance to herbivores and a need to explore indirect effects via recruitment of natural enemies in greater detail.

Variable effects across chewing and sucking herbivores were observed with rhizobacterial inoculation of host plants more strongly reducing chewing insect behaviours (host choice and feeding) than fitness traits, and sucking herbivores primarily impacted through reduced reproduction (fecundity), but also reduced body sizes and life span. This is likely a reflection of the feeding behaviour. Sucking insects feed on plant sap using modified tube-like mouthparts causing less mechanical damage to plant tissues than chewing herbivores (Walling 2000). This mode of feeding by sucking insects means they may by-pass some of the plant’s immediate defences triggered by tissue damage, but may be more susceptible to other induced defences following sap-ingestion (Züst & Agrawal 2016). In contrast, chewing insects disrupt plant tissues more extensively, triggering induction of rhizobacteria-primed broad-spectrum plant defences. These may be more immediate and thus influence host-choice and feeding rate before any detectable effect on growth or reproductive output.

There were strong differences across the rhizobacteria species, with *Bacillus* spp (Blake, Christensen & Kovács 2021) inoculants providing stronger general effects than the commonly studied *Pseudomonas* spp (Pieterse *et al*. 2021). We grouped all other rhizobacteria species since alone they had insufficient data, but they also significantly impacted insects across various traits. These effects were not restricted to specific insect or plant families also highlighting the generalisability of rhizobacterial inoculation for plant protection (Biere & Bennett 2013; Gruden *et al*. 2020). While most chewing herbivores studied were Lepidopterans, we observed significant behaviour and fitness effects of rhizobacteria on Chrysomelidae leaf beetles and variation among Lepidopteran sub-families. Across host plants, *Bacillus* inoculations were most effective against chewing herbivores for both Brassicaceae and Poaceae plants with limited effect of *Pseudomonas* inoculants. Aphids were the most studied sucking herbivores, most affected by *Pseudomonas* spp and ‘other’ rhizobacteria, many of which belonged to nitrogen-fixing rhizobia species. This may indicate an interplay between plants and microbes that can modulate nutrient availability and induce plant defences, i.e. as biostimulants (plant-growth-promoting rhizobacteria) and bioprotectants (priming and induction of defence pathways).

Single-strain inoculants generally performed better than multi-strain or community inoculations (stronger reduction of insect traits). However, this may be due to the choice of multi-strain inoculant communities as opposed to loss of effect. Interestingly, for the sucking herbivores, single-strain *Bacillus* inoculants performed better than multi-strain *Bacillus* inoculants, indicating potential strain incompatibility could affect outcomes (Díaz *et al*. 2023). The reduced abundance of *Bacillus* strains in the rhizosphere over time following inoculation (Moshe *et al*. 2024) may also impact variable effectiveness of a multi-strain community, particularly if they use similar resources that diminish over time. This may also explain the increased effectiveness of multi-genus inoculants for reducing sucking insect fitness on inoculated plants, as observed in the meta-analysis results. The potential of diverse community inoculants (also termed SynComs, synthetic communities) to provide a broad array of benefits is high, and as such is a keen focus of current plant microbiome research (Pascale *et al*. 2020; Marin, Gonzalez & Poupin 2021; Shayanthan, Ordoñez & Oresnik 2022; Fagorzi *et al*. 2023; Poudel *et al*. 2023; He *et al*. 2024; Wang *et al*. 2024). Strain and functional compatibility will be key to enable SynComs to persist and deliver benefits to the host plants over time (De Souza, Armanhi & Arruda 2020; O’Banion *et al*. 2020; Song *et al*. 2020; Hayashi, Fujita & Toju 2024).

The majority of experimental work is conducted in environmentally-controlled conditions, often using sterilise soil in sterilised pots and watered with sterilised water. This is a good approach to uncover the effect of a focal inoculated strain, but it has been difficult to translate to the field which is messy and diverse. Our results showed that while the overall effects were significant for pot-based experiments but not for the field trials, there were several field trials reporting highly significant outcomes. A limitation of this data is that it was restricted to host choice (only on *Bacillus* spp inoculated plants) for chewing herbivores, and fecundity (averaged across all microbes, but mostly *Bacillus* spp or rhizobia) for sucking herbivores. Nevertheless, the increased native microbiota of field soils and environmental variability will reduce the ability of inoculants to colonise and persist among plant roots (Gruden *et al*. 2020; Frew *et al*. 2022; He *et al*. 2024).

In conclusion, there is great potential for using rhizobacteria to increase plant resistance to insect pests but these meta-analysis results show that we need to consider the broader interacting environment when developing effective inoculants (Song *et al*. 2020; Shayanthan, Ordoñez & Oresnik 2022). This will involve understanding which rhizobacteria are important for specific plants, against specific insects or under specific environments, but also those that provide broad benefits across multiple ecological systems. With this knowledge, we can develop synthetic microbial communities with broad acting functions across multiple plants, crops and environments.

## Acknowledgements

This work was supported by a BBSRC (UKRI) David Phillips Fellowship BB/S010556/1 to SEZ, BBSRC NLD studentship to MP, and German Research Council, Deutsche Forschungsgemeinschaft (DFG; project number 397565003) funding to SZ and OSM.

## Notes

### Competing Interest Statement

The authors have declared no competing interest.

### Summary of Updates

Figures updated, introduction and discussion revised.

